# A Potential Novel COVID-19 Vaccine With RBD-HR1/HR2 Hexamer Structure

**DOI:** 10.1101/2021.10.28.465226

**Authors:** Hongbo Liu, Xiang Gao, Guoyong Wang, Jianjun Zhang, Jiajie Zhou, Tingting Wei, Yu Zhang, Yujiao Liu, Jinhua Piao, Qiulei Zhang, Yayuan Wang, Xin Ma, Xiaoting Zhu, Yikun Rao, Wenjuan Xia, Heng Xie, Wei Zhang

## Abstract

The COVID-19 pandemic and the continued spreading of the SARS-CoV-2 variants have brought a grave public health consequence and severely devastated the global economy with recessions. Vaccination is considered as one of the most promising and efficient methods to end the COVID-19 pandemic and mitigate the disease conditions if infected. Although a few vaccines have been developed with an unprecedented speed, scientists around the world are continuing pursuing the best possible vaccines with innovations. Comparing to the expensive mRNA vaccines and attenuated/inactivated SARS-CoV-2 vaccines, recombinant protein vaccines have certain advantages, including their safety (non-virus components), potential stronger immunogenicity, broader protection, ease of scaling-up production, reduced cost, etc. In this study, we reported a novel COVID-19 vaccine generated with RBD-HR1/HR2 hexamer that was creatively fused with the RBD domain and heptad repeat 1 (HR1) or heptad repeat 2 (HR2) to form a dumbbell-shaped hexamer to target the spike S1 subunit. The novel hexamer COVID-19 vaccine induced high titers of neutralizing antibody in mouse studies (>100,000), and further experiments also showed that the vaccine also induced an alternative antibody to the HR1 region, which probably alleviated the drop of immunogenicity from the frequent mutations of SARS-CoV-2.

## INTRODUCTION

The COVID-19 pandemic and the continued spreading of the SARS-CoV-2 variants have brought a grave public health consequence and severely devastated the global economy with recessions. By the end of August, 2021, more than 220 million infections were diagnosed with COVID-19 and 4.5 million people died of this deadly disease. The pandemic has overwhelmed the most health care systems and depleted tremendous life-saving medical resources around the world, adversely impacting on not only the COVID-19 patients but also other patient populations with various diseases and timely-sensitive medical conditions, including patients in numerous clinical treatment trials.

Vaccination is considered as one of the most promising and efficient methods to end the COVID-19 pandemic and mitigate the disease conditions if infected. So far, a few vaccines have been developed with an unprecedented speed originated from the close collaboration among pharmaceutical companies, academic centers, government regulatory agencies, *etc*. These vaccines have been developed with different technologies such as novel mRNA, DNA, viral vector-based complex, recombinant protein, and the traditional attenuated/inactivated SARS-CoV-2 virus^1^. The different vaccines all have preventive effects with different safety, efficacy and cost-effective profiles.

Scientists around the world are continuing pursuing the best possible vaccines with innovations, aiming to improve current vaccine’s safety, efficacy, cost-effectiveness, accessibility, etc. to help address the public health challenges.

Comparing to the expensive mRNA vaccines and attenuated/inactivated SARS-CoV-2 vaccines, recombinant protein vaccines have certain advantages, including their safety (non-virus components), potential stronger immunogenicity, broader protection, ease of scaling-up production, reduced cost, *etc*^1,2^. According to the published studies, the SARS-CoV-2 spike protein is the major component of capsid proteins and most neutralizing antibodies have been developed to target the spike protein to block the virus’ entering into the host cells by interfering with the binding and interaction between receptor-binding domain (RBD) of the spike S1 subunit and the receptor of angiotensin-converting enzyme 2 (ACE2) of the host cell^2–4^. In this study, we reported a novel COVID-19 vaccine generated with RBD-HR1/HR2 hexamer that was creatively fused with the RBD domain and heptad repeat 1 (HR1) or heptad repeat 2 (HR2) to form a dumbbell-shaped hexamer to target the spike S1 subunit. The novel hexamer COVID-19 vaccine induced high titers of neutralizing antibody in mouse studies, and further experiments also showed that the vaccine also induced an alternative antibody to the HR1 region, which probably alleviated the drop of immunogenicity from the frequent mutations of SARS-CoV-2.

## MATERIALS AND METHODS

### Protein expression and purification

The constructs of RBD (319-550) and fusion proteins were synthesized and sub-cloned into pZD vector between EcoRI and HindIII. Expi293 was used for transient expression of all RBD proteins. Cytiva Q Sepharose FastFlow was used for protein captures and the proteins were further purified by gel filtration to remove contaminants. The final purified proteins were analyzed by SDS-PAGE and LC-MS. For SEC-HPLC analysis, around 100 μg protein was loaded onto Superdex 200 Increase using PBS as running buffer.

### In vitro binding assay

HR1 with C-terminal poly-his tag was immobilized onto a NTA sensor chip (from GE Healthcare) to around 500 RU using a Biacore SPR 8K (GE Healthcare) and a running buffer composed of PBS buffer and 0.05% Tween-20 was used. Serial dilutions of the HR2 peptide (synthesized by Genscript) were flowed through with a concentration ranging from 0.05 to 100 nM and the data were fit to a 1:1 binding model using Biacore Evaluation Software (GE Healthcare).

### Negative-staining EM study

Protein samples (5 μL) were applied to TEM grids (Ultra-Thin Carbon Coated Grids, Beijing XXBR Technology) for approximate 1 min and wicked away by filter paper. Addition of 5 μL of deionized water was then applied and wicked away immediately. A solution of 1% uranyl acetate (5 μL) was placed on the grid for 1 min, wicked away, and air-dried. TEM was performed using a Talos L120C transmission electron microscope (accelerating voltage 120 keV) at a magnification of 57kx and 92kx, respectively. The calculated pixel size is 0.152nm at 92kx. The total electron dose is 30e-/Å^2^.

### Pseudotyped-virus neutralization assay

Mice were divided into seven groups with 8 female mice in each group and alhydrogel adjuvant (1.25 mg/ml, InvivoGen) was used for the animal study. In this study, the final protein concentration was diluted into 0.1 mg/ml after mixed with the adjuvant and administrated at 10 μL per mice (BALB/c). The antigen was administrated 3 times on day 1, 15 and 29, the antibody titers for RBD domain were continuously monitored. In pseudotyped-virus neutralization experiments, the pseudotyped-virus was constructed based on pseudotyped ΔG-luciferase (G*ΔG-luciferase) rVSV with SARS-CoV-2 spike protein gene inserted. Three types of pseudotyped-virus, the original SARS-CoV-2, Delta (B.1.617.2) and Lambda (C.37) strains were tested with several serial dilutions. Bright-Lumi^™^ kit was used for fluorescence detection and the final IC50 (50% neutralization) was calculated on GraphPad Prism.

## RESULTS AND DISCUSSION

In contrast to the RBD domain of S1 subunit, both of the HR1 region and the HR2 region are from the S2 subunit and are considered as the key components for virus entering host cells. HR1 forms a homo-trimeric parallel alpha-helix structure, which exposes three highly conserved hydrophobic grooves on the surface. Each HR2 monomer lays into the grooves to assemble into an antiparallel six-helix bundle hexamer core structure (Fig.1a), mainly by hydrophobic interactions^5^. We thus synthesized the HR1 and HR2 peptides and examined the binding affinity of SARS-CoV-2 HR1 and HR2 by Biacore-SPR 8K. The data indicated that the binding affinity between HR1 and HR2 were strong (dissociation constant K_D_=2.24 nM, Fig.1b). Previous studies showed that the oligomerization of RBD increased its immunogenicity and thus improved the titers of neutralizing antibody. The stable native HR1/HR2 hexamer led us to think if we could use this property to generate a HR1/HR2-RBD hexamer as a novel design for improved COVID-19 vaccines (Fig.1a), we then constructed a RBD-HR1 and RBD-HR2 fusion protein respectively. As there was a long flexible region C-terminal to RBD^6^, we didn’t introduce any linker sequence between RBD and HR1 or HR2. The final two constructs were composed of pure virus sequences without any artificial sequence. The two proteins were expressed in HEK293 cells and purified by chromatography method. LC-MS analyses showed that all disulfide bonds were formed and the proteins were well-folded. In order to further verify the function of RBD-HR1 trimer, we tested the binding between RBD-HR1 and its binding receptor ACE2 by Fortebio (Fig.s1). In this experiment, the RBD-HR1 showed a very similar binding affinity to ACE2 as RBD itself did, indicating the formation of well-folded RBD-HR1 protein. We further mixed the purified RBD-HR1 and RBD-HR2 proteins with stoichiometric ratio 1 to 1. Both HPLC-SEC and electron microscope (EM, negative staining) studies showed that RBD-HR1 and RBD-HR2 formed a stable hexamer in the solution (Fig.1c and Fig.1d). The EM study clearly demonstrated that the RBD-HR1/HR2 hexamer exhibited a dumbbell-shaped structure, in which the six RBD domains were connected by a stable and long HR1/HR2 alpha-helix bundle. This unique shape potentially mimicked the trimeric structure of SARS-CoV-2 spike protein and exposed more epitopes from both RBD and HR regions, leading to an increase in immunogenicity.

**Fig.1.**
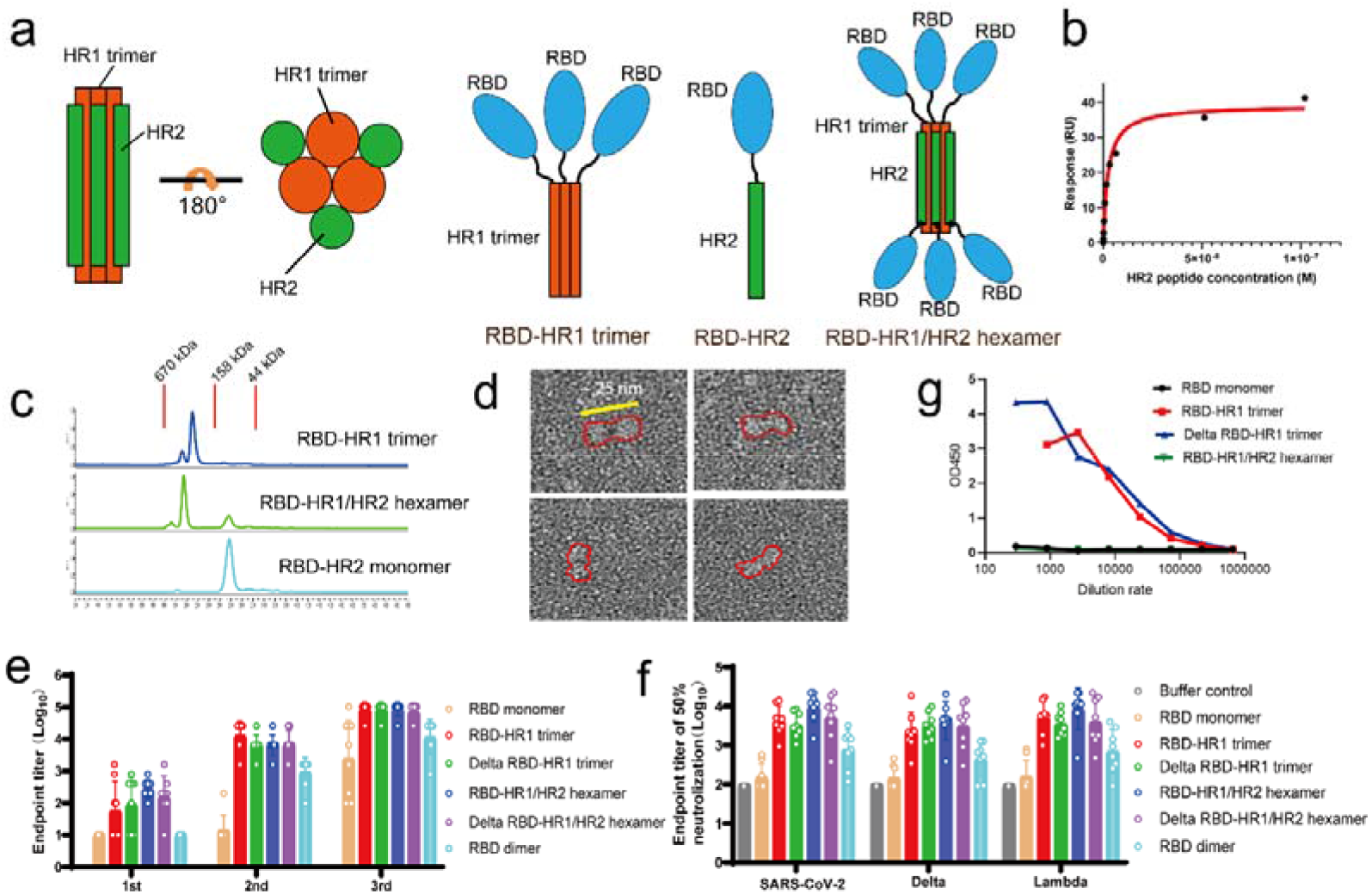
Schematic illustration of RBD-HR1/HR2 hexamer and pseudotyped-virus neutralization assay. **(a)** Schematic structure of RBD-HR1 or -HR2 fusion protein. The RBD-HR1 and RBD-HR2 could form a stable hexamer through the tight interaction between HR1 and HR2. (**b)** SPR binding test for HR1 (immobilized in this assay) and HR2. (**c)** SEC-HPLC analysis of RBD-HR1/HR2 hexamer. (**d)** The dumbbell-shaped RBD-HR1/HR2 hexamer under electron microscope (negative-staining). (**e)** Both RBD-HR1 trimer and RBD-HR1/HR2 hexamer could generate high anti-RBD titers in the animal study. (**f**) The serum from RBD-HR1 trimer or RBD-HR1/HR2 hexamer immunized mice efficiently neutralize different types of SARS-CoV-2 pseudotyped-viruses. (**g)** RBD-HR1 trimer could induce high titer of anti-HR1 antibody.

In order to test and compare the immunogenicity of this hexamer protein with other reported forms of RBD, we constructed and purified the RBD monomer (319-550, D614G stain), and a dimeric RBD^7^ and also the RBD domain from the Delta strain (B.1.617.2). Alhydrogel adjuvant (1.25 mg/ml, InvivoGen) was used for the animal study, and the final protein concentration was diluted into 0.1 mg/ml and administrated at 10 μl per female mouse (BALB/c). The antigen was administrated 3 times on day 1, 15 and 29, the antibody titers for RBD domain were continuously monitored. In our experiment, both trimeric RBD-HR1 and hexametric RBD-HR1/HR2 induced very high antibody titers (above 100,000) which was almost 10-fold of the reported RBD dimer and 1000-fold of the RBD monomer (Fig.1e). Statistical analysis didn’t show significant difference between the common RBD and the one from the Delta strain, indicating the hexamer might be used as a novel vaccine development strategy for COVID-19 and its variants. In pseudotyped-virus neutralization experiments, we tested the serum samples from the vaccinated mice with three types of pseudotyped-viruses, the original SARS-CoV-2, Delta (B.1.617.2) and Lambda (C.37) strains, to investigate whether the antigen generated antibody could potentially protect the mice from different SARS-CoV-2 strains (Fig.1f). Our results showed that the same mouse serum neutralized viruses from all three virus strains and could generate the same degree of protection among them, indicating the vaccine we developed might be potentially effective not only for the original SARS-CoV-2 but also for other mutants of SARS-CoV-2 strain.

So far, RBD remains the main epitope for most vaccine development. However, RBD is one of the most unstable regions and mutates rapidly, thus makes effective vaccine either short-lived or less effective during the pandemic, leading to more virus infections and disease spreading. HR1 and HR2 have been reported to present more epitopes that could generate more alternative neutralizing antibodies to inhibit SARS-CoV-2 infections^8,9^. For example, in our trimeric RBD-HR1 (Fig.1a), the HR regions were exposed and could generate antibody. Subsequently, we measured whether there was any anti-HR1 or anti-HR2 antibody in the mouse serum. The mouse serum was analyzed by Biacore SPR with HR1 or HR2 and immobilized onto CM5 chip (Fig.s2). Our data showed that the trimeric RBD-HR1 induced anti-HR1 antibody, while no anti-HR1 antibody was detected from hexametric RBD-HR1/HR2 immunized mouse. This was probably due to the HR2 region that shield part of the epitope from HR1. No anti-HR2 antibody was detected for all analyzed serum samples. Last, we also randomly selected two batches of serum samples from RBD-HR1 immunized mice and detected the titer of anti-HR1 antibody. Our data showed that the titer of anti-HR1 antibody (~ 50,000) was about half of anti-RBD antibody (~100,000) in average, but the level of 50,000 was still relatively high and offered protection (Fig.1g).

In summary, the COVID-19 pandemic and the newly emerged Delta and Lambda strains, continue to be the greatest global public health threat. More efficient and cost-effective vaccines with a broader spectrum of prevention and protection for various SARS-CoV-2 variants are in great need for coping with the current public health crisis and global economic recovery^10^. We reported a potential novel COVID-19 vaccine with RBD and HR1/HR2. The vaccine induced high titer of anti-RBD neutralizing antibody in mice and provided more and broader neutralizing antibodies against the different virus strains. Similarly, the trimeric RBD-HR1 formation also induced an alternative anti-HR1 neutralizing antibody, which probably mitigated the spreading of the fast-mutating COVID-19 in mice. As hexametric HR1/HR2 complex formation is the general mechanism of class I enveloped virus, this RBD-HR1/HR2 hexamer design may be applied to other virus vaccine development and offers a potential novel strategy to develop improved human vaccines. Our novel COVID-19 vaccine with RBD-HR1/HR2 hexamer structure may offer a great potential for COVID-19 and its various variants.

## ACKNOWLEDGEMENTS

We thank Shuimu BioSciences for help with the negative–staining EM and Joinn biologics for the pseudotyped–virus neutralization assay.

## AUTHOR CONTRIBUTIONS

H. L., X. G., W. X., and W. Z. conceived this study. All authors carried out or designed the experiments and manuscript preparation. H. X. helped to revise the whole paper.

## ADDITIONAL INFORMATION

The experimental methods and some additional data could be found in the supplementary information.

