## Supplementary information for "A Potential Novel COVID-19 Vaccine With RBD-HR1/HR2 Hexamer Structure"

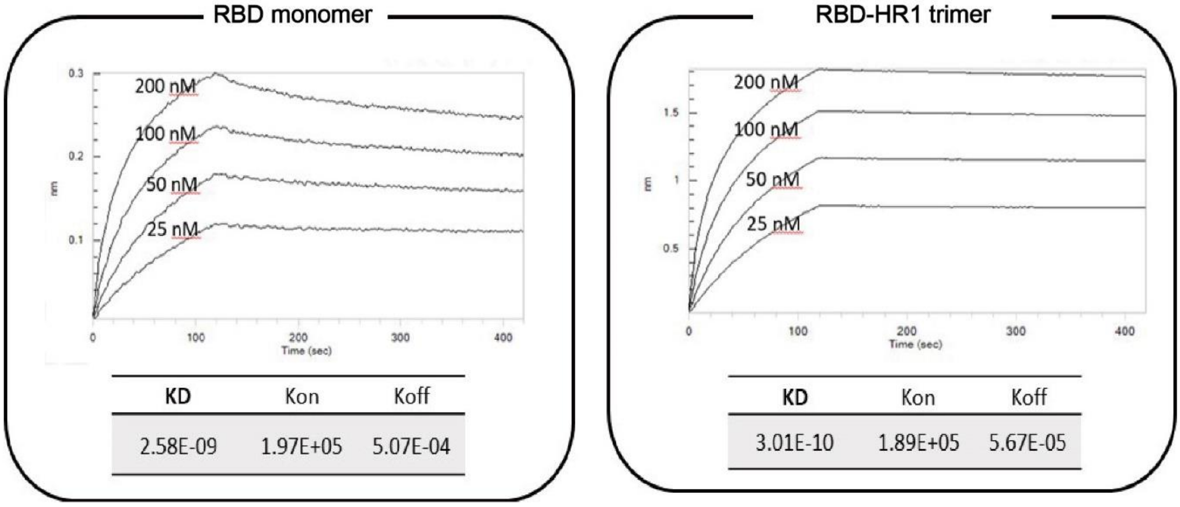


Fig. S1 In vitro binding test for RBD monomer and RBD-HR1 trimer to ACE2 receptor by Fortebio. The RBD or RBD-HR1 trimer was immobilized onto AR2G Biosensor and the analyte ACE2 was titrated from 25 nM to 200 nM.

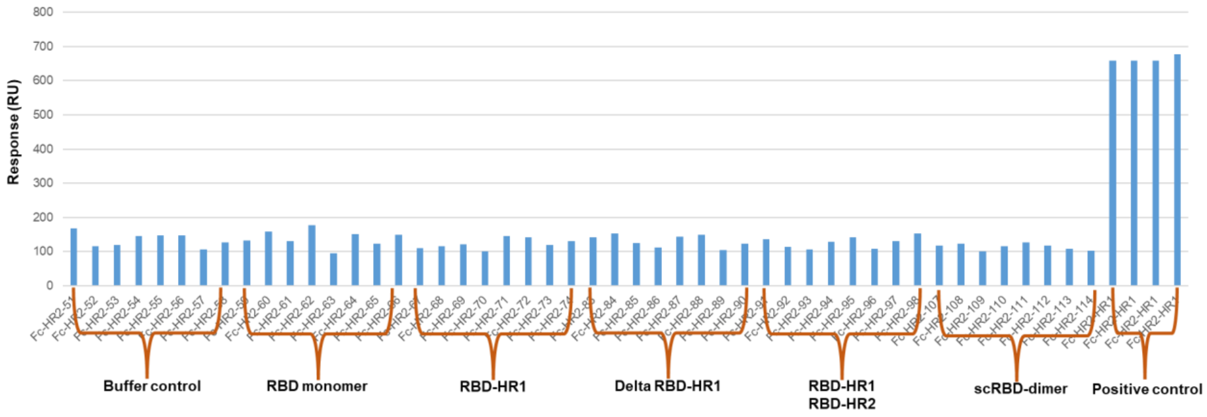


Fig. S2 Detection of anti-HR1 antibodies from the immunized mouse serum. In this assay, HR1 was immobilized onto CM5 sensor chip and different immunized mouse serum was flowing through the chip with contacting time of 250 seconds. The response units at the end of contact time were collected for this analysis.
